# Sentence-level neural dynamics in speech motor cortex

**DOI:** 10.64898/2026.11.10.765804

**Authors:** H. Levi-Aharoni, J. J. Jude, S. Haro, A. J. Acosta, S. B. Allcroft, M. C. Ottenhoff, N. S. Card, M. Wairagkar, C. Iacobacci, A. L. Pritchard, B. Richards, D. M. Brandman, S. D. Stavisky, N. Au Yong, C. Pandarinath, A. C. Paulk, P. N. Hadar, J. Li, Z. M. Williams, S. S. Cash, J. D. Simeral, L. R. Hochberg, D. B. Rubin

## Abstract

Prior recordings from speech motor cortex have revealed phonemic, syllabic, articulatory and speech-sequence representations, while more recent studies indicate that activity may also reflect higher-order features of speech production. However, whether speech motor cortex is sensitive to the linguistic organization of sentences remains unclear. Here we show that single-unit activity in speech motor cortex contains information about sentence-level linguistic structure. In three BrainGate2 participants with implanted microelectrode arrays in the left precentral gyrus, we recorded neural activity during an instructed-delay speech task comprising English sentences, word lists, syntactically structured nonword sentences and nonword lists. Neural activity in areas 6v and 55b contained speech- and language-related components, with stronger stimulus-condition effects in 55b, and stimulus condition was decodable in all three participants. In two larger datasets, semantic categories of English sentences were decodable from neural activity. These findings demonstrate that speech motor cortex encodes information about linguistic structure and semantic content at the level of sentences.

## Introduction

Speech production requires the transformation of linguistic representations into precise sequences of articulatory motor commands. Intracortical recordings from speech motor cortex have revealed detailed representations of phonemes and articulatory movements, supporting increasingly sophisticated brain-computer interfaces (BCIs) for communication. Recent demonstrations of implantable BCIs (iBCIs) for speech restoration have inferred intended speech by decoding acoustic features of attempted speech (Wairagkar et al. 2025; Silva et al. 2024; Herff et al. 2019; Anumanchipalli et al. 2019) or sequences of attempted phonemes (Willett et al. 2023; Metzger et al. 2023; Card et al. 2024; Jude et al. 2026; Nason-Tomaszewski et al. 2026) directly from neural activity in motor cortex. Predicted sequences can then be passed through a language model to reconstruct the most likely words and phrases (Willett et al. 2023). While this approach can effectively restore communication for users (Card et al. 2024, 2026), variability in phoneme decoding accuracy across users persists (Willett et al. 2023; Metzger et al. 2023; Jude et al. 2026; Nason-Tomaszewski et al. 2026). Understanding what linguistic information is present in speech motor cortex beyond these relatively low-level representations could both inform models of speech production and identify additional signals for speech neuroprostheses.

Recent studies have shown that activity in speech motor cortex reflects increasingly complex aspects of speech production, including the sequencing of speech sounds and word-level structure (Silva et al. 2022; Liu et al. 2025; Xu et al. 2025; Gadonneix et al. 2026; Krasa et al. 2026). These findings also suggest that speech motor cortex is engaged during the preparation of upcoming speech, with activity varying according to the complexity and stage of the planned speech sequence (Silva et al. 2022; Liu et al. 2025; Xu et al. 2025). However, whether speech motor cortex contains sentence-level linguistic information, beyond the lower-level speech representations that comprise sentences, remains unknown (Yeaton 2025).

To test whether speech motor cortex contains sentence-level linguistic information beyond phonemic content, we used an instructed-delay copy task in which participants attempted to read aloud visually presented text-based stimuli that varied in linguistic content. In addition to two English sentence datasets (Card et al. 2024; Jude et al. 2026) based on the Switchboard corpus (Godfrey et al. 1992), we used stimuli drawn from four conditions: English sentences, English word lists, Jabberwocky sentences, and nonword lists (pronounceable pseudowords) (Fedorenko 2010), commonly used in neuroimaging studies of sentence structure and lexical-semantic sensitivity (Fedorenko et al. 2016, 2020; Shain et al. 2024; Pylkkänen 2019; Matchin and Hickok 2020). We used this task in research sessions conducted with three participants currently enrolled in the BrainGate2 clinical trial (ClinicalTrials.gov Identifier: NCT00912041). All three participants have impaired speech production due to paralysis (participants T15 and T17 due to amyotrophic lateral sclerosis (ALS) and participant T16 due to brainstem stroke) and underwent placement, guided by the Human Connectome Project (HCP) parcellation pipeline, of four to six 64-channel NeuroPort microelectrode arrays in the left precentral gyrus, including in speech motor cortex: T17 and T16 in areas 6v and 55b; T15 in areas 6v, 55b, and 4 (Fig. 1b).

**Figure 1:**
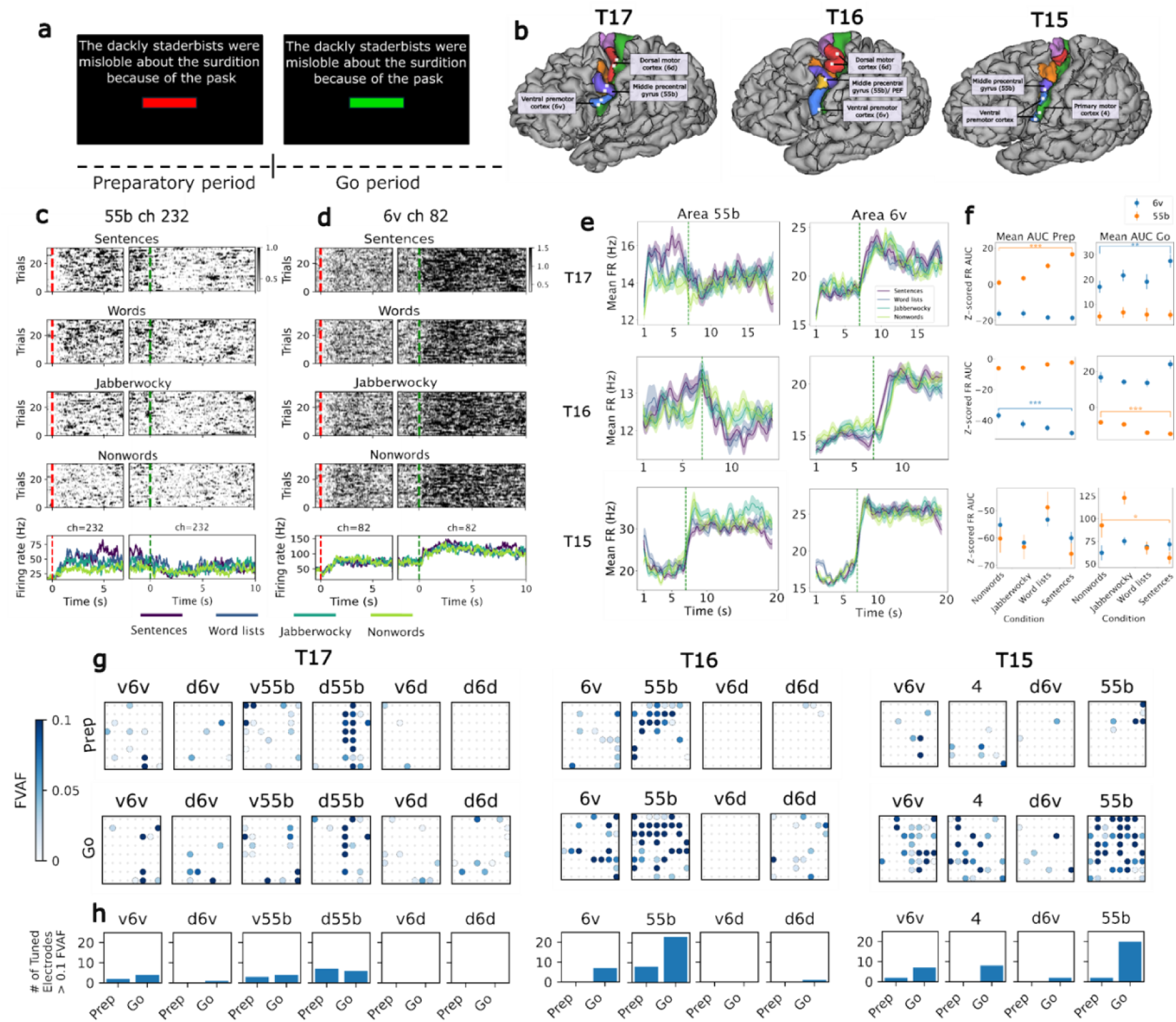
Tuning to different linguistic task conditions across motor cortex areas in an instructed delay copy task. **a)** The instructed delay copy task: during the preparatory period, text was presented on the screen above a red rectangle. The participant was instructed to read the text and prepare to attempt to speak it. After a brief variable delay (7.0 ± 0.5 s), the rectangle turned green and a beep sounded, which signaled the participant to attempt to speak aloud the text displayed on the screen. **b)** Microelectrode array locations for T17, T16 and T15. All arrays are located in the language-dominant left hemisphere. For T17, two arrays were placed in each of the following cortical areas: dorsal (6d), middle (55b) and ventral (6v) precentral gyrus. For T16, two arrays were placed in dorsal (6d), one in middle (55b) and one in ventral (6v) precentral gyrus. For T15, one array was placed in middle (55b), two in ventral (6v) and one in posterior (4) precentral gyrus. Brain images adapted from (Deo et al. 2026) with the author’s permission. **c)** Example raster plots ordered by condition from an illustrative channel in area 55b from participant T17. Bottom panel: averaged peristimulus time histogram (PSTH) per condition for the corresponding channel, smoothed using a 0.1 s convolution window. The panels are showing the trials aligned to trial onset (left, marked by red dashed vertical line) and to go cue (right, marked by green dashed vertical line). **d)** Same as c, for an illustrative channel in area 6v. **e)** Averaged PSTH for area 55b (left) and area 6v (right) of each participant. Firing rates were averaged over all 128, 64, and 64 channels in each area for participants T17 [v6v+d6v, v55b + d55b], T16 [6v, 55b], and T15 [v6v, 55b] respectively. For visual clarity, the x-axis runs from 1 second after stimulus onset through stimulus offset (a large stimulus onset response at t=0 otherwise obscures task specific responses; for the full response see Extended Data Figure 4). **f)** Mean area under the curve (AUC) of the channel-averaged z-scored single-trial firing rate (FR) for preparatory period (left) and go period (right) for each of the areas (blue: 6v, orange: 55b). Error bars represent standard error of the mean (SEM). Asterisks denote significant differences between the nonwords and the sentences conditions (*: p<0.05, **: p<0.01, *** p<0.001, Student’s T-test). **g)** Tuning heatmaps for all arrays of each participant. Circles indicate that threshold crossing features on a given electrode varied significantly across all conditions during the prep period (top) and during the go period (bottom) (p < 0.05, assessed with one-way analysis of variance). Shading indicates the fraction of variance accounted for (FVAF) by cross condition differences in threshold crossing rate. **h)** Number of tuned electrodes where threshold crossing features on a given electrode varied significantly across all conditions, separated by task period (p < 0.05 assessed with one-way analysis of variance).

Across all three participants, we found evidence that speech motor cortex contains information about linguistic structure at the level of entire sentences. Neural responses recorded from both areas 6v and 55b were tuned to stimulus condition during both preparation and attempted production of speech. Although the strength of this tuning varied across participants, it was overall stronger in area 55b than in area 6v. Moreover, neural activity from both cortical areas was predictive of the semantic category of intended sentences in two larger datasets collected with two of the participants. Across these complementary analyses, we found evidence that speech motor cortex contains information about sentence-level linguistic structure and semantic content.

Together, these findings show that as linguistic content is transformed into an articulatory motor program, decodable semantic information is present in speech motor cortex, revealing a potentially valuable signal for improving iBCI decoding of intended speech.

## Results

### Areas 6v and 55b are differentially tuned for articulation versus language during speech preparation and production

To assess for differences in tuning to language across areas of speech motor cortex, participants performed a language-localizer task adopted from the fMRI literature (Fedorenko 2010). In an instructed delay paradigm, participants attempted to read aloud strings of visually presented text randomly drawn from one of four stimulus classes. These stimuli included (1) grammatically correct sentences, (2) word lists, (3) Jabberwocky (intact sentence structure with nonsense content words), and (4) nonword lists (Fig. 1a).

We observed prominent area-related differences in responses in participant T17. Electrode arrays targeting Brodmann area 6v, which is thought to contain a core articulatory control region of motor cortex (Willett et al. 2023; Card et al. 2024), captured robust firing rate responses to all four stimulus classes (Fig. 1d). Neural responses recorded from 6v were most prominently following the Go cue, sustained throughout the trial, and showed minimal modulation to stimulus condition (Fig. 1e). In contrast, neural activity from area 55b was more prominent during the preparatory period, and firing rates appeared more substantially modulated by stimulus condition (Fig. 1c, 1e).

To quantify these differences observed in participant T17’s arrays, we calculated the average area under the firing rate curve (AUC) for each trial in the channel-averaged activity. For participant T17, neural activity in area 6v increased significantly more during the Go period than during the preparation period, whereas area 55b displayed the opposite behavior (Fig. 1f and Extended Data Fig. 6). Moreover, the overall neural response recorded from area 55b, but not area 6v, varied significantly (p < 0.001) across stimulus condition during the preparatory period. In contrast, the neural response recorded from area 6v, but not area 55b, varied significantly (p < 0.01) across stimulus condition during the Go period (Fig. 1f). Interestingly, in this participant, the preparatory activity in area 55b and the Go period activity in area 6v both appeared sensitive to the level of semantic content of the stimulus conditions, with the most consistently robust response elicited by grammatically intact English sentences, smaller responses to unstructured word lists and Jabberwocky, and relatively minimal signal modulation for unstructured lists of nonwords (Fig. 1c, 1f).

For participant T16, we observed a different pattern of responses. As in participant T17, we observed significantly greater increases in firing rates in area 55b during the preparatory period than during the Go period, and significantly increased firing rates in area 6v during the Go period compared to the preparatory period (Fig. 1f, Extended Data Fig. 6, p < 0.001. However, unlike in T17, firing rates in area 6v were significantly different between the nonwords and English sentences conditions during the preparatory period (p < 0.001) but not the Go period, and responses recorded from area 55b showed a similar stimulus class modulation during the Go period (p < 0.001) but not during the preparatory period (Fig 1f). For participant T15, we observed a significant difference in the AUC between the nonwords and English sentences conditions only in 55b during the Go period.

While these averaged response profiles varied considerably across participants, in all three participants, the fraction of variance in stimulus response accounted for (FVAF, see Methods) by activity in area 55b was consistently greater than that accounted for by activity recorded from area 6v (Fig. 1g). Similarly, the number of electrodes significantly tuned to stimulus condition was greater in 55b than 6v (Fig. 1h). These findings suggest that area 6v may play a greater role in encoding articulatory motor processes, whereas area 55b may be more involved in upstream, language-specific processing.

### Dynamical structure reveals differences in motor- and language-related activity in areas 6v and 55b

We next sought to better understand how information pertaining to stimulus condition is represented across cortical areas. As a first assessment, we quantified the decoding accuracy for stimulus condition for each participant, during both the preparatory and Go periods, using neural activity from each cortical region (Fig. 2a,b,c. Also see Extended Data Fig. 1). For participant T17, stimulus condition decoding accuracy during the preparatory period was significantly higher in 55b than in 6v (p < 10^-5^). For participant T16, stimulus condition decoding was also greater in 55b than in 6v (p <0.001), however, this difference was observed during the Go period rather than the preparatory period. For participant T15, decoding accuracy using neural signals from ventral 6v (v6v) was higher than from 55b; this was also observed during the Go period (p <0.01).

**Figure 2:**
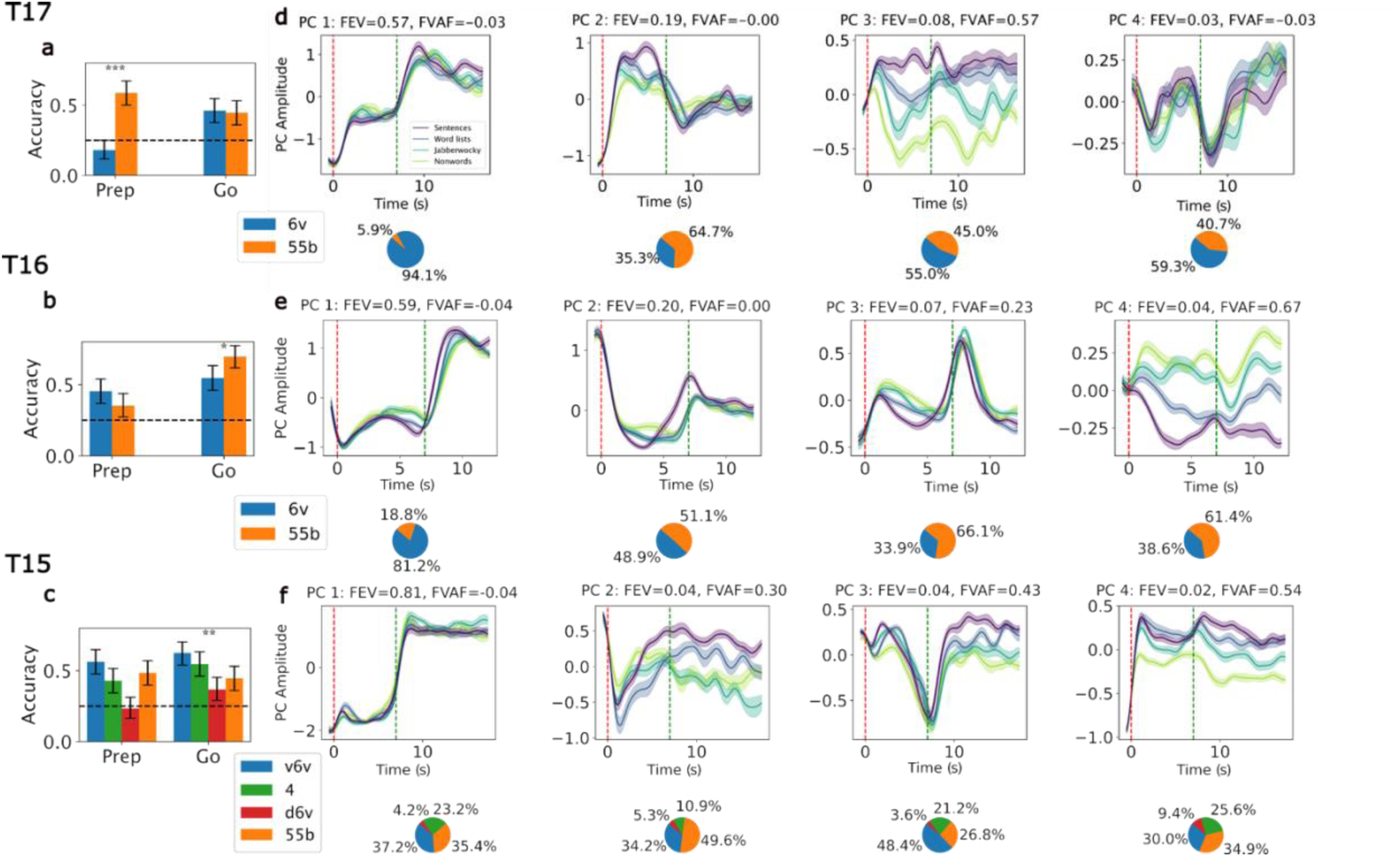
Areas 6v and 55b show different separability of task condition. **a,b,c)** Stimulus condition decoding accuracy using a Gaussian Naive Bayes (GNB) classifier in each cortical area, separated into preparatory and attempted speech (Go) periods, using only the first 6.5 seconds for both Prep and Go. Asterisks denote significant difference between the 6v and 55b decoding accuracies. Error bars represent 95% confidence interval estimated using 10,000 bootstrap samples. Also see Extended Data Fig. 1. **d,e,f)** PCA waveforms of the first four PC components, averaged over all trials of each condition for each participant. Dashed vertical lines denote cue onset (red) and averaged Go cue time (7 s, in green. The pie chart under each plot shows the contribution of activity from each cortical area to the PC weights, quantified as the sum of squared weights from channels belonging to each area.

To further characterize the neural representation of stimulus condition and to better understand the observed differences in response profiles between the participants, we applied principal component analysis (PCA) to the trial-averaged neural activity. Across all participants, PCA revealed two broad classes of components: (i) components aligned to the temporal structure of motor activation (i.e., with a large difference in activity between preparation and attempted articulation) with minimal sensitivity to stimulus condition, and (ii) components strongly modulated by the stimulus condition. Notably, for participants T17 and T16, who showed the most prominent differences between the responses of 55b and 6v, the first PC showed minimal condition dependent modulation (FVAF<0 for both, meaning the condition-averaged mean was a worse predictor of the trial firing rate than the mean over all trials) and had minimal contribution from 55b (also see Extended Data Fig. 2).

As we had observed in the firing rate PSTH from participant T17 (Fig. 1e), the PCs that were most modulated by stimulus condition in each participant (i.e. PC3 for T17, and PC4 for T16 and T15) exhibited the same lexical-semantic ordering in the magnitude of responses for the four stimulus conditions (Fig. 2d,e,f). This recurring pattern observed across participants, despite differences in array location and participant clinical phenotype, suggests a common representation for these levels of linguistic content. Overall, although raw firing rate averages varied considerably across participants and cortical areas, these PCA results suggest that the dominant low-dimensional projections of cortical responses share three key characteristics across participants: (i) a dichotomy between motor-dominant and language-dominant patterns of activity, (ii) a greater contribution from area 55b over area 6v neurons to language-dominant patterns, and (iii) within language-dominant patterns, a preserved ordinal ranking of lexical content across the four stimulus classes.

### Low-dimensional trajectories reveal sentence-level condition-dependent dynamics

To explore the temporal dynamics of responses, we used the first three PCA components to display the averaged trajectories for each condition in the low-dimensional PCA space. These trajectories showed a similar rotational pattern that varied in shape across conditions and participants (Fig. 3a). Interestingly, for T17 we observed that the sentences and word lists stimulus conditions appeared to trace smoother trajectories in state space than the Jabberwocky and nonwords stimulus conditions (Fig. 3a, left). Previous work on natural language processing and visual perception have proposed that neural networks (both artificial and biological) may optimize predictions by learning to straighten neural trajectories in representation space (Hénaff et al. 2019, 2021; Hosseini and Fedorenko 2023). To test this hypothesis and to quantify these differences in trajectories across conditions and participants, we calculated the curvature along the trial trajectories in the first four PCs (see Methods). Consistent with this hypothesis, during the beginning of the Go period, all participants showed a higher curvature (i.e., less straightened trajectory) for the nonwords compared to the sentences condition (Fig. 3b, c).

**Figure 3:**
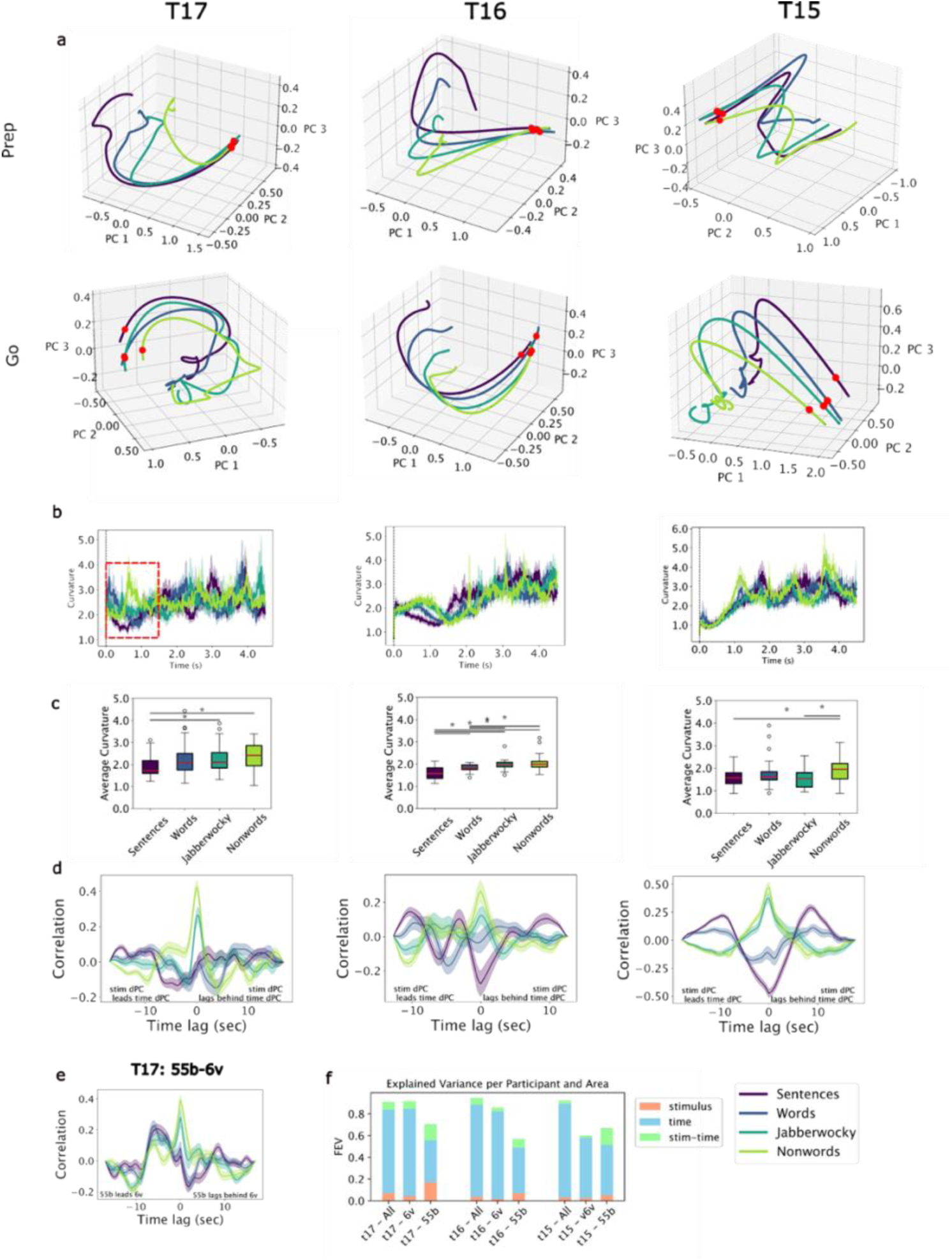
Dynamical response characteristics to different task conditions. **a)** Condition-averaged PCA trajectories in the top 3 PCA components space. Top: preparatory period. Red dot marks stimulus onset. Bottom: Go period. Red dot marks the go cue. **b)** Trajectory curvature along the first four PCA dimensions, shown for the first 4.5 seconds of the Go period. **c)** Mean curvature over the initial 1.5 seconds of the Go period (marked in dashed red line on panel b, left). Asterisks mark significant difference (p<0.05). **d)** Cross-correlation in time between the first stimulus dPCA component and the first time dPCA component. Decomposition was performed over the entire trial duration. Stability was assessed via 1,000 random half-splits (Pearson’s r = 0.90 ± 0.12 for T15, 0.58 ± 0.30 for T16, and 0.68 ± 0.18 for T17) **e)** Cross-correlation in time between T17’s 55b channel-averaged signal and 6v channel-averaged signal. **f)** Fraction of explained variance (FEV) in All channels, 6v channels (v6v for T15) and 55b channels by the first four dPCA components of each of the marginalizations (stimulus, time and stimulus-time).

As previously described, the PCA projection of each participant’s neural activity revealed a mix of PCs that were more informative of either motor activation or stimulus condition. To more specifically probe the relative contribution of these two factors to the overall activity, we employed demixed PCA (dPCA) (Kobak 2016), which can extract and differentiate time-dominant and stimulus-dominant components from neural population activity. The time-dominant component here is taken to represent the Go-cue-triggered initiation of motor related activity (i.e., articulation), whereas the stimulus-dominant component captures the variance related to stimulus condition (i.e., language). Using this approach, we find that stimulus-dominant dPCA components explained more of the variance in 55b channels than in 6v channels (Fig. 3f).

Using this dPCA representation, we next explored the interaction between the time-dominant activity and the condition-dominant activity by examining the time-lagged cross-correlation between the first stimulus and time components (Fig. 3d, Extended Data Fig. 5). Across all participants, we observed a more similar correlation pattern between the Nonwords and Jabberwocky conditions compared to other pairs (Pearson’s r = 0.56, 0.78 and 0.95 for T17, T16 and T15 respectively. Also see Extended Data Fig. 7). In addition, for participants T16 and T17 we observed a negative peak in negative lags for English sentences and Word lists, suggesting an inverse relationship between language and speech component activity, with a temporal offset between the two. For comparison, we also included the 55b-6v time-lagged correlation plots for T17 (Fig. 3e).

### Population activity in speech motor cortex encodes semantic information

Lastly, we asked whether the representational structure observed in the prior analyses could be extended to semantic distinctions between categories of natural English sentences. We analyzed two large datasets of attempted speech of English sentences drawn from the Switchboard corpus (Godfrey et al. 1992): one collected with T17 (Jude et al. 2026) and one with T15 (Card et al. 2024) and used only trials that were at least 6 seconds in duration (1864 sentences for T17, 5358 sentences for T15). Sentences were embedded using a pretrained BERT [[Sentence-BERT: Sentence Embeddings using Siamese BERT-Networks](https://huggingface.co/papers/1908.10084)] model and were clustered using k-means (see Methods), yielding 7 clusters for T17 and 10 clusters for T15 (see Methods for details). Trial counts were equalized across clusters by trimming to the lower trials count across clusters (100 and 134 trials per cluster for T17 and T15 respectively). The silhouette scores of these two clustered datasets are 0.049 for T17 and 0.046 for T15. This means that despite pruning there was still a considerable overlap between classes, posing a challenging classification task with the neural data.

We visualized the averaged trajectories for each cluster on the first stimulus dPC, which revealed moderate cluster-level separation (Fig. 4c,f). We then trained a Gaussian Naive Bayes (GNB) classifier to decode cluster identity from the recorded neural activity. Neural activity was averaged across the trial period for each channel, and for T17, we excluded the two arrays placed in area 6d. For T17, the decoding accuracy was 23% when using signals only from area 55b, 20% for signals from 6v, and 24% when using both areas (chance is 14%). For T15, decoding accuracy was 17% for both 55b and v6v separately and 20% for all 256 channels in all four arrays (chance is 10%). While in T17’s dataset decoding accuracy was higher in 55b than in 6v, paralleling the pattern observed in the language localizer task, in T15’s dataset the accuracies were comparable across 55b and v6v (Fig. 4e), consistent with the overall similarity between T15’s 6v and 55b response properties. We also tested the decoding accuracy using only the first four stimulus dPCs, resulting in 25% accuracy for T17 and 16% for T15, suggesting that this low dimensional subspace captures most of the relevant semantic information within the neural activity of areas 6v and 55b, at least as measured by GNB decoding accuracy.

**Figure 4:**
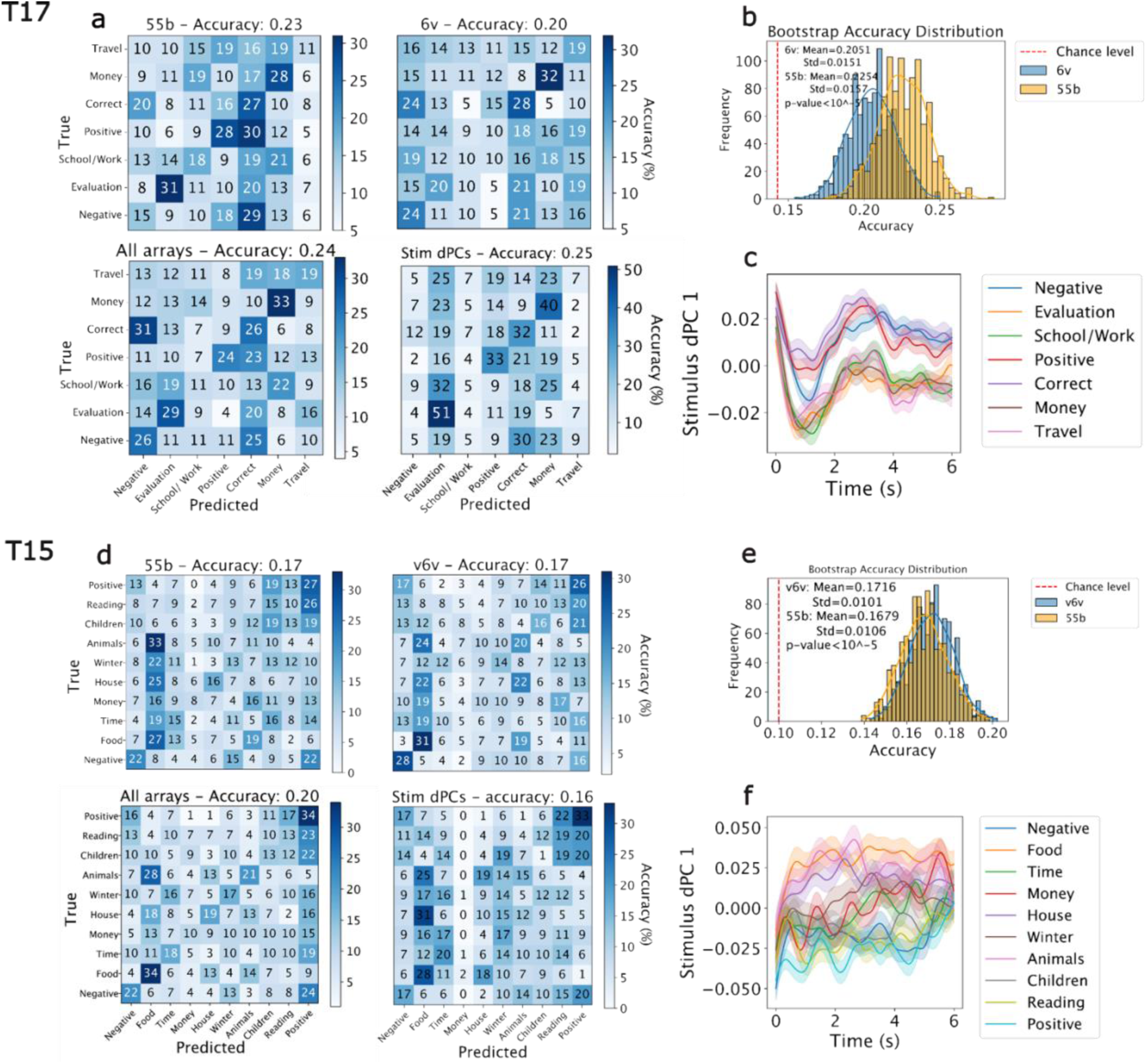
Classification of semantic categories in Areas 6v and 55b. **a)** Confusion matrices for 7 semantic classes extracted using unsupervised clustering from T17’s corpus of sentences, calculated using a GNB decoder applied on the first 6 seconds after trial onset, shown for Area 55b (128 channels) (top left), Area 6v (128 channels) (top right), all channels (55b and 6v – 256 channels) (bottom left), and using the first 4 dPCA stimulus components that were computed on held out trials not used for the classification training (see Methods for details). Cluster topics were extracted manually based on unsupervised clustering of the dataset sentences (see Methods and Extended Data). **b)** Distribution of classification accuracy obtained via 1000 bootstrap resampling. Red dashed line indicates chance level. **c)** The trial-averaged dPCA activity of the first stimulus component for each of the semantic classes. **d)** Confusion matrices for 10 semantic classes extracted using unsupervised clustering from T15’s corpus of sentences, calculated using a GNB decoder applied on the first 6 seconds after trial onset (regardless of Go-cue timing), shown for Area 55b (64 channels) (top left), Area v6v (64 channels) (top right), all channels (v6v, d6v, 55b and area 4, 256 channels) (bottom left), and using the first 4 dPCA stimulus components. **e)** Same as (b). **f)** Same as (c).

## Discussion

In this work we provide evidence that sentence-level structure is represented within human speech motor cortex, specifically in areas 6v and 55b, regions until recently understood as primarily articulatory (6v) (Mugler et al. 2014; Pei et al. 2011; Anumanchipalli et al. 2019; Willett et al. 2023) and syllabic (55b) (Silva et al. 2022; Liu et al. 2025) in function. Linguistic structure was reliably reflected in the neural population dynamics of both areas: activity systematically varied as stimulus condition differed in semantic content, and both regions were sensitive to stimulus condition during both preparation and attempted speech. These effects were expressed within cortical areas classically associated with speech execution, demonstrating that sentence-level information is present in the neural dynamics supporting speech production.

Sentence-level structure was represented across ventral (6v) and middle (55b) precentral gyrus in a distributed yet biased manner. Although neurons in area 6v were more strongly engaged during attempted speech and in area 55b during preparation (with a greater proportion of the variance in activity recorded from area 55b accounted for by stimulus class), activity in both regions encoded both linguistic and articulatory information about upcoming speech. Thus, rather than a strict division between articulatory and linguistic computations, these results support a model of graded specialization, in which networks in both cortical areas participate in processing higher-order linguistic structure.

We further explored differences in neural dynamics across conditions by quantifying the curvature of low-dimensional population trajectories. Although linguistic surprisal, and more specifically probability of words, was not directly manipulated in this study, stimuli with higher linguistic regularities were associated with smoother trajectories in the initial Go period phase, whereas unfamiliar stimuli exhibited more sharply curved trajectories. This pattern has an intriguing correspondence with a recent theoretical proposal that linguistic predictability shapes the geometry of neural trajectories in large language models (Hosseini and Fedorenko 2023). In this framework, smoother trajectories reflect more predictable and resource-efficient dynamics, which potentially require fewer inter-area control signals, whereas sharply curved trajectories may necessitate tighter coupling and increased information exchange between regions. This interpretation aligns with the higher zero-lag correlations observed for unfamiliar stimuli (nonwords and word lists in Fig. 3e) and suggests that semantic content influences neural coordination across areas during speech production.

Some of the motivation for this study is that a growing understanding of language representation in the human brain will continue to reveal opportunities for restoring neurologic function impaired by injury or disease. To that end, it is notable that sentence-level activations supported above-chance semantic classification in speech BCI training datasets in both ventral (area 6v) and mid (area 55b) precentral gyrus. For the two switchboard datasets (from T17 and T15), semantic categories could be decoded from neural activity in each region, indicating that sentence-level semantic information is distributed across the speech motor system. The top four projections obtained with demixed PCA preserved all or most of the information relevant for this semantic category decoding, suggesting that sentence-level semantic information is represented in low-dimensional population activity in speech motor cortex.

Several limitations should be noted. First, our focus on language production within a BCI framework means that reading-related activity is interpreted as preparatory, and the present study does not allow us to dissociate perceptual processes from preparatory activity. Future work will further investigate preparatory vs. perceptual processes including in more naturalistic communication (Cai et al. 2025; Castellucci et al. 2022; Kunz et al. 2025; Cai et al. 2026), word-level surprisal (Hou et al. 2025) and different types of linguistic predictability and their relationship to neural trajectory geometry (Hosseini and Fedorenko 2023; Yan et al. 2026). In addition, several factors could have contributed to the differences in the neural responses to each condition observed in the language localizer task. Future work will be aimed at dissociating the contribution of semantics versus syntax, task difficulty and working memory load. While areas 6v and 55b may be considered motor areas and hence not affected by these factors, the fact that there is lexical-semantic information affecting the dynamics warrants a closer look at the effect of these other cognitive processes.

Second, this study was conducted in three participants with intact language but marked or complete speech impairment, which may have influenced neural activity in speech motor cortex. The presence of substantial similarities across participants, including between an individual with complete speech motor paralysis due to advanced ALS and an individual with stroke-related speech impairment, suggests that the core findings reported may reflect consistent underlying biology that persists despite neurologic impairment.

Finally, while our participant cohort exhibited consistent properties in area 6v, greater variability was observed in area 55b. Possible contributors to the observed heterogeneity may include individual differences in functional organization and in particular in relation to handedness (T15 is left-handed while T17 and T16 are right-handed), possible variability within area 55b and the stochastic nature of neural sampling inherent in the use of microelectrode arrays. Lastly, the process by which functional parcellation was performed, using the MRI-based Human Connectome Project approach, similarly may introduce variability in the Brodmann area boundaries due to noise in the MRI acquisition process and subtle variability across clinical sites in other aspects of the imaging analysis pipeline. More broadly, while many studies explored the relationship of fMRI to other, more refined electrophysiological measurements (e.g. Siero et al. 2014) the relationship between surface-projected cortical parcellation maps and more highly resolved, cortical layer-specific neuronal ensemble recording is unknown (Hermosillo et al. 2024; Zaldivar et al. 2022; Ojemann et al. 2013; Hermes and Siero 2022). While analyzing neural activity in a low-dimensional space partially mitigated some of these challenges in the current study, larger participant cohort will be required to more precisely delineate the unique contribution of area 55b to language processing and production.

In summary, these findings place area 55b within the core language-to-speech production network, where sentence-level linguistic structure is transformed into motor plans through structured interactions with area 6v. While area 6v remains more strongly biased toward articulatory execution, both regions express sentence-dependent dynamics that reflect the global linguistic structure. This organization supports a modular but interactive architecture in which linguistic and motor representations are only partially segregated, and are dynamically coupled during speech production, refining existing models of language production and revealing extensive integration between language and motor systems.

## Methods

### Participants and array implantation

All research sessions were conducted with participants enrolled in the BrainGate2 clinical trial (ClinicalTrials.gov ID: NCT00912041, Registration Date: 2009-06-01, CAUTION: Investigational Device. Limited by Federal Law to Investigational Use). All procedures were approved by the relevant institutional review boards (at Mass General Brigham for all participants, at Brown University and the VA Providence Healthcare System for T17, at Emory University for T16, and at University of California, Davis for T15), and informed consent was obtained from all participants. Participant-specific MRI-based localization procedures have been published previously (Deo et al. 2026; Card et al. 2024; Nason-Tomaszewski et al. 2026; Hadar et al. 2026; Jude et al. 2026). All research sessions were performed at the participants’ place of residence.

Participant T17 is a 34-year-old (at the time of data collection) right-handed man with ALS. He has tetraplegia, anarthria, and ventilator dependence. At the time of this study, his only remaining volitional motor control was over his extraocular muscles. He has six 64-channel intracortical microelectrode arrays (NeuroPort/MultiPort, Blackrock Microsystems; 1.5 mm electrode length) placed in the left hemisphere, guided by preoperative anatomical MRI, task-based functional MRI, and cortical parcellation using the Human Connectome Project (HCP) multi-modal parcellation pipeline (Hadar et al. 2026). Two arrays were placed in the HCP-designated dorsal precentral gyrus (Brodmann area 6d), two arrays were placed in the ventral precentral gyrus (Brodmann area 6v), and two arrays were placed in middle precentral gyrus (Brodmann area 55b). The language localizer task was performed by T17 about 2.5 months post device placement (trial day 83) and the open loop sentences dataset was collected over a period of several months (trial days 46-89) as described in detail in (Jude et al. 2026).

Participant T16 is a 52-year-old right-handed woman with a history of tetraplegia and dysarthria due to a brainstem stroke that occurred 19 years prior to enrolling in the study. She underwent pre-operative anatomical and functional MRI and has four microelectrode arrays (NeuroPort/MultiPort, Blackrock Microsystems; 1.5 mm electrode length) placed in the left precentral gyrus; two targeting area 6d, one targeting area 6v, and one targeting area 55b. Of note, functional assessments reported elsewhere suggest that the array targeting area 55b may be in the border between 55b and FEF (Nason-Tomaszewski et al. 2026). For further details on pre- and post-operative array localization for T16 see (Nason-Tomaszewski et al. 2026). The language localizer task was performed by T16 about 15.5 months post implantation (trial day 466).

Participant T15 is a 45-year-old left-handed man with ALS who had symptoms that began five years prior to enrollment in this study. At the time of this study, T15 was non-ambulatory, had dysarthric speech and no functional use of his hands, and used an eye-tracking system as his primary means of communication. Based on preoperative anatomical MRI and HCP cortical parcellation, four 64-channel intracortical microelectrode arrays (NeuroPort/MultiPort, Blackrock Microsystems; 1.5 mm electrode length) were placed in the left (language-dominant) mid- and ventral precentral gyrus (two targeting Brodmann area 6v, one targeting area 55b, and one targeting area 4). Notably, T15’s two arrays in area 6v had markedly different response properties to a number of experimental paradigms, likely owing to their substantial anatomic distance, and so, in contrast to participant T17, activity recorded from these arrays was not combined to report an overall average activity for the area. The language localizer task was performed by T15 about 13.5 months post implantation (trial day 410) and the open loop sentences dataset was collected over a period of several months as described in detail in (Card et al. 2024) (trial days 25-636).

### Tasks and stimuli

#### Task 1: Language localizer task

Participants were asked, in an instructed delay paradigm, to produce speech from one of four classes of cues; English sentence, lists of English words, grammatically structured sentences with content words replaced by nonwords (“Jabberwocky”), or nonword lists (Fedorenko 2010). Data was collected from each participant in a single session. The experimental task consisted of four blocks, each lasting approximately 15 minutes. Each text cue consisted of 12 “words”; cue types were randomly ordered within each block, with each block containing eight cues from each type, resulting in 32 trials per block with the same cue files used across all participants. Each trial began with a preparatory delay period of 7 ± 0.5 seconds (uniformly jittered within this range), during which the cue was displayed on the computer screen above a red rectangle. At the end of this delay, the rectangle turned green and a beep sounded, signaling the go cue for the participant to begin attempting to speak the sentence. Each trial was ended manually by the participant; T16 and T17 made an eye movement to the researcher to signify their having completed attempting to speak; participant T15 used an eye-gaze tracking device to signify his completion of each trial. Go-period duration varied across participants and was truncated for analysis to the minimal go period duration across trials (which resulted in 16.9 s, 12.7 s, or 17.8 s for T17, T16, and T15, respectively).

#### Task 2: Instructed delay sentence production task

For the sentence-level analysis (T17 and T15), English sentences from the Switchboard corpus (Godfrey et al. 1992) were used in a similar instructed delay copy task structure as in Task 1. The delay period for this task was 4.5 seconds, and the go period was determined similarly as for task 1: using an operator button press triggered by an eye gaze with T17 and using an eye tracker system with T15. Full details about these datasets can be found in (Jude et al. 2026) for T17 and in (Card et al. 2024) for T15.

### Neural signal acquisition and preprocessing

Neural voltage time series signals were recorded and digitized (30 kHz, 16 bits per sample) using the Neuroport system (Blackrock Neurotech). Neural data were streamed in real time from the neural signal processor (NSP) to our signal processing pipeline, which was run in a custom-built software suite (Ali et al. 2024). Signals were first analog filtered (T17, T15: band-pass filter, 250 Hz to 5 kHz, acausal 4th order Butterworth filter. T16: high-pass filter, 250 Hz, acausal 4th order Butterworth filter) using the Scipy python library (scipy.signal.filtfilt). Subsequently, linear regression referencing (LRR) was performed using the filtered digitized data recorded during an initial reference block performed at the beginning of each session. LRR coefficients are computed by solving *Y* = *W X* where *Y* is the signal from a given electrode we require and *X* is the signal from all other electrodes. We solve for the LRR weight matrix through least squares calculation *W*: *W* = *inv*(*X^T^ X*)*X^T^ Y* where *inv* is matrix inversion. Electrode-specific thresholds were calculated to identify spike events. Thresholds were set at -3.5 times the standard deviation of the voltage signal per electrode (i.e. -3.5 RMS) for T17, and at -4.5 RMS for T16 and T15. The number of non-causal threshold crossings (ncTX) were computed by counting the number of times the filtered neural time series crossed these calculated thresholds (threshold crossing rates). Additionally, spike band power was computed by taking the sum of squared voltages observed during each 10 ms time bin.

To account for neural non-stationarities (drifts in mean firing rate), for each electrode, threshold crossing rates and spike band power were Z-scored (mean subtracted and divided by standard deviation per electrode for each block), except for raster plots. Feature extraction (threshold crossings and spike band power), binning, decoding and task phase control were performed through the Python based, modular BRAND framework (Ali et al. 2024), where each process is instantiated as a self-contained node-based Python program. Messaging between these nodes is performed using a Redis database. In post-session offline analysis we performed additional artifact removal steps: (1) removing large magnitude artefacts by zeroing all bins larger than 10 standard deviations from the mean, (2) smoothing the signal using a Gaussian kernel applied to windows of N=100 bins with \sigma=N/6. These additional steps were performed for all analyses except for the decoding presented in Fig. 2a,b,c.

### Condition tuning and fraction of variance accounted for (FVAF)

Condition-dependent tuning for each channel was quantified using the fraction of variance accounted for (FVAF) by across-condition differences in mean firing rate: *FVAF* = 1 − *SS_Err_*/*SS_Tot_*, where *SS_Tot_* is the total sum of squared errors from the mean across all trials and *SS_Err_* is the total sum of squared errors from the condition mean. FVAF values were computed separately for preparation and go epochs. Significance was assessed with one-way analysis of variance. Channels with p<0.05 were considered significantly modulated by the stimulus condition.

Note that in this context the FVAF is a measure of stimulus-dependent modulation and can be calculated both for channels and components. This is different from the fraction of explained variance (FEV) that is calculated here for PCA and dPCA components, and is a measure of the variance explained by components (regardless of the stimulus), defined by 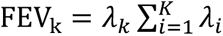 for the kth component from a total of K components, with *λ_k_* being the kth eigenvalue.

### Correlation analyses

We computed per-trial cross-correlation matrices and then averaged over all trials per condition to construct a correlation matrix per condition. The correlation SEM per pair was calculated over all trials per condition. Sentence minus nonword difference matrices were constructed to visualize condition-dependent changes; negative values reflect higher correlation in nonword trials. Bar plots summarizing these differences across participants are provided in Extended Data Fig. 3.

### Principal component analysis

PCA was applied to trial-averaged neural activity concatenated across conditions. Single-trial data was then projected onto the learned PC space. PCs were defined per participant and sorted by variance explained. Area contributions to each PC were quantified as the sum of squared weights from channels belonging to each area. To obtain the relative contribution per area, each area’s contribution was normalized by the sum of contributions of all areas. Condition-related modulation of each PC was quantified using fraction of variance accounted for (FVAF), paralleling the single-channel analysis.

### Demixed PCA (dPCA)

To extract condition- and time-specific low-dimensional structure, we performed demixed PCA (dPCA) (Kobak et al. 2016) with stimulus and time marginalizations using dPCA python package (Machens Lab, https://github.com/machenslab/dPCA). For the semantic decoding analysis, to avoid information leakage, dPCA was computed only on a training subset of trials (50%, separate from the classification training set), and the resulting transformations were applied to the held-out 50% test data (on which a GNB leave-one-out classification was performed). This procedure was repeated across 1000 bootstrap samples.

### Trajectory curvature estimation

For each trial, instantaneous curvature was approximated by first computing velocity vectors from consecutive trajectory points, normalizing these to unit tangent vectors T^(t), then taking the differences of these unit tangents. This discrete approximation corresponds to the standard differential geometry definition of curvature: κ(t) = ||dT^/ds||, where s denotes arc length. This yields a time-resolved curvature profile for each trial. For visualization purposes, the displayed curvature time course was smoothed using \sigma = 6.67ms (1/3 bins)

### Sentence-embedding clustering

Sentence embeddings were generated for each participant’s sentence database (T17: 1864 sentences, T15: 5358 sentences) using the *bert-base-nli-mean-tokens* model from the Sentence Transformers library ([Sentence-BERT: Sentence Embeddings using Siamese BERT-Networks](https://huggingface.co/papers/1908.10084)). The clustering analysis was intended as a proof of concept for semantic sentence clustering. Accordingly, the analysis was restricted to a limited number of a relatively well-separated clusters within each database, extracted using the procedures detailed below.

After defining the clusters, trial counts were equalized by truncating each cluster to the minimum number of trials across clusters, yielding 100 trials per cluster for T17 and 134 trials per cluster for T15.

To avoid information leakage from the dPCA analysis, the decomposition was computed only on the training set in each bootstrap iteration (90% of trials), then applied to the held-out 10%, which was classified using leave-one-out cross-validation. This procedure was repeated 100 times.

#### T17 dataset sentence clustering

For participant T17, the procedure consisted of the following steps. First, k-means clustering (k = 50) was applied to the sentence embedding space. From these, 30 clusters exhibiting the lowest variance in a two-dimensional PCA projection were selected, and sentences belonging to all other clusters were discarded. The remaining sentences were then re-clustered into 20 clusters. From these, 10 clusters with maximally separated centroids were selected, as determined using `sklearn’s pairwise_distances_argmin_min`. Outlier sentences were subsequently removed by discarding sentences whose distance from their assigned centroid exceeded the 90th percentile. This final set of seven centroids was used to assign cluster labels to the full set of sentences, followed by an additional round of outlier rejection applied to the complete dataset. To ensure sufficient data per cluster, clusters containing fewer than 100 sentences were discarded.

#### T15 dataset sentence clustering

For T15, the larger dataset allowed us to simplify the cluster extraction procedure: k-means clustering (k = 50) was first applied to the sentence embedding space. Next, 15 clusters were randomly selected, and their silhouette score was computed. This random selection was repeated 1,000 times, and the set of 15 clusters with the highest silhouette score was retained (score = 0.137). Then, five clusters with ambiguous or poorly defined topics were manually discarded based on the 20 sentences closest to each cluster centroid, resulting in 10 well-defined clusters.

### Gaussian Naive Bayes

The Naive Bayes classifier is a simple probabilistic predictor which assumes conditional independence between features given the class. Gaussian Naive Bayes takes this a step further for continuous data and makes the additional assumption that values associated with each class are normally distributed. Single-trial classification results were generated using a cross-validated (leave-one-out) Gaussian Naive Bayes classifier. All trials are z-scored (mean subtracted and standard deviation divided) and time-averaged in the go or preparatory period of each task. For every trial, the classifier is trained on all other trials and evaluated on the held-out trial. We use the Scikit-learn 54 implementation of Gaussian Naive Bayes.

### Sentence category decoding

For both stimulus condition and semantic clusters, decoding was performed using a Gaussian Naive Bayes (GNB) classifier applied to either (i) channel activity, or (ii) projections onto the first four stimulus dPCs. For both cases the classifier was applied on the averaged activity per channel/component in the specified period (e.g. Go period). For accuracy estimate with the dPCA components, dPCA was computed on 50% of trials, test-set projections were computed on the remaining 50%, and classification performance was evaluated using leave-one-out cross-validation. This procedure was repeated 100 times. The same GNB classifier and leave-one-out scheme were used for decoding the four text categories of the language localizer task.

### Data availability

All data required to reproduce the findings in this study will be made publicly available upon publication. Switchboard datasets are publicly available from (Jude et al. 2026; Card et al. 2024). The dataset contains intracortical neural signals recorded from participants along with detailed information about the BCI behavior tasks and performance metrics.

### Code availability

The code for reproducing the figures will be made available upon publication.

## Acknowledgements

We thank T17, T16, T15 and their families and carepartners for the time and effort they contributed to the BG2 trial. We thank Maryam Masood and David Rosler for expert regulatory contributions enabling this research.

## Funding Statement

This work was supported by Office of Research and Development, Department of Veterans Affairs (N2864C, A4820R, A2295R, A3803R); NIH NIDCD (U01DC017844, U01DC019430, K23DC021297, 1DP2DC021055); NIH NINDS (DP2NS127291), AHA (23SCEFIA1156586); CDMRP (HT94252310153), Brown University Provost’s STEM Postdoctoral Fellowship to S. Haro; A.P. Giannini Postdoctoral Fellowship to N. Card.

## Competing interests

The MGH Translational Research Center has a clinical research support agreement (CRSA) with Ability Neuro, Axoft, Neuralink, Neurobionics, Precision Neuro, Synchron, and Reach Neuro, for which LRH provides consultative input, and for Paradromics, for which LRH and DBR provide consultative input. LRH is a non-compensated member of the Board of Directors of a nonprofit assistive communication device technology foundation (Speak Your Mind Foundation). Mass General Brigham (MGB) is convening the Implantable Brain-Computer Interface Collaborative Community (iBCI-CC); charitable gift agreements to MGB, including those received to date from Paradromics, Synchron, Precision Neuro, Neuralink, and Blackrock Neurotech, support the iBCI-CC, for which LRH provides effort. Stavisky is an inventor on intellectual property owned by Stanford University that has been licensed to Blackrock Neurotech and Neuralink Corp. Wairagkar, Card, Stavisky, and Brandman have patent applications related to speech BCI owned by the Regents of the University of California. Stavisky is an advisor to Subsense and was a consultant to Neuralink. Brandman was an ad-hoc consultant for Paradromics Inc. at the time of data collection. He is an ad-hoc consultant for Globus Medical. Pandarinath is a research scientist at Meta (Reality Labs).

CAUTION: Investigational Device. Limited by Federal Law to Investigational Use. The content is solely the responsibility of the authors and does not necessarily represent the official views of the National Institutes of Health, or the Department of Veterans Affairs, or the United States Government.

## Extended data

**Extended Data Figure 1.**
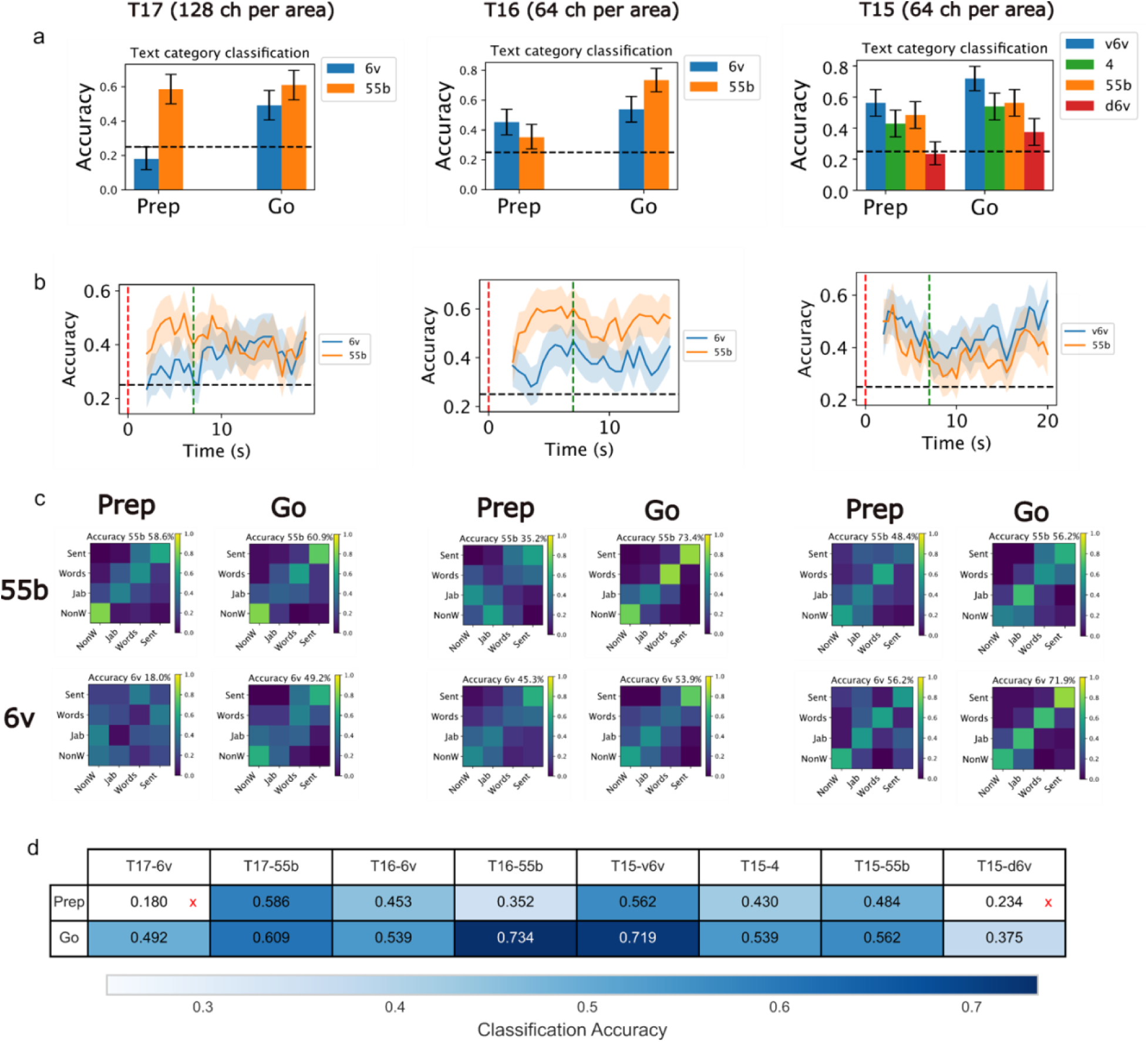
Gaussian Naïve Bayes decoding performance in the language localizer task. **a)** Stimulus condition decoding accuracy in each cortical area for each participant, separated by preparatory (Prep) and attempted speech (Go) periods using the first 6.5 s for the Prep period and the minimum epoch duration across all trials for the Go period. Error bars represent 95% confidence interval estimated using 10,000 bootstrap samples. **b)** Time-resolved GNB category decoding accuracy for areas 6v and 55b. Each point shows the decoding accuracy using a 2-second window ending at that time point. Accuracy was calculated at 0.5-second intervals. Shading shows the 95% confidence interval estimated using 10,000 bootstrap samples. **c)** Confusion matrices for GNB category classification, separated by preparatory (Prep) and Go periods, for areas 6v (v6v for T15) and 55b. The title of each matrix shows the mean accuracy. **d)** GNB decoding accuracy values. Values are corresponding to the bar plots in a. A red x denotes a non-significant value based on the confidence interval value.

**Extended Data Figure 2:**
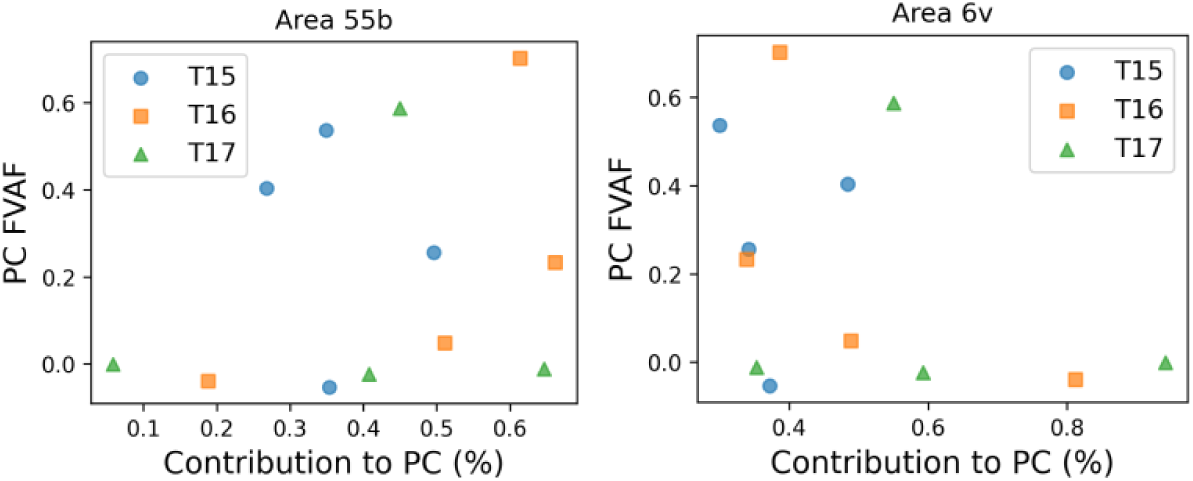
Principal components FVAF as a function of area contribution. For each principal component (PC) as defined in Figure 2 in the main text, the PC’s fraction of variance accounted for (FVAF) is shown as a function of **(left)** area 55b’s and **(right)** area 6v’s (or v6v for T15) contribution to that PC. Contribution is defined as the root sum of squared coefficients across all channels in a given area, as in Figure 2 in the main text.

**Extended Data Figure 3:**
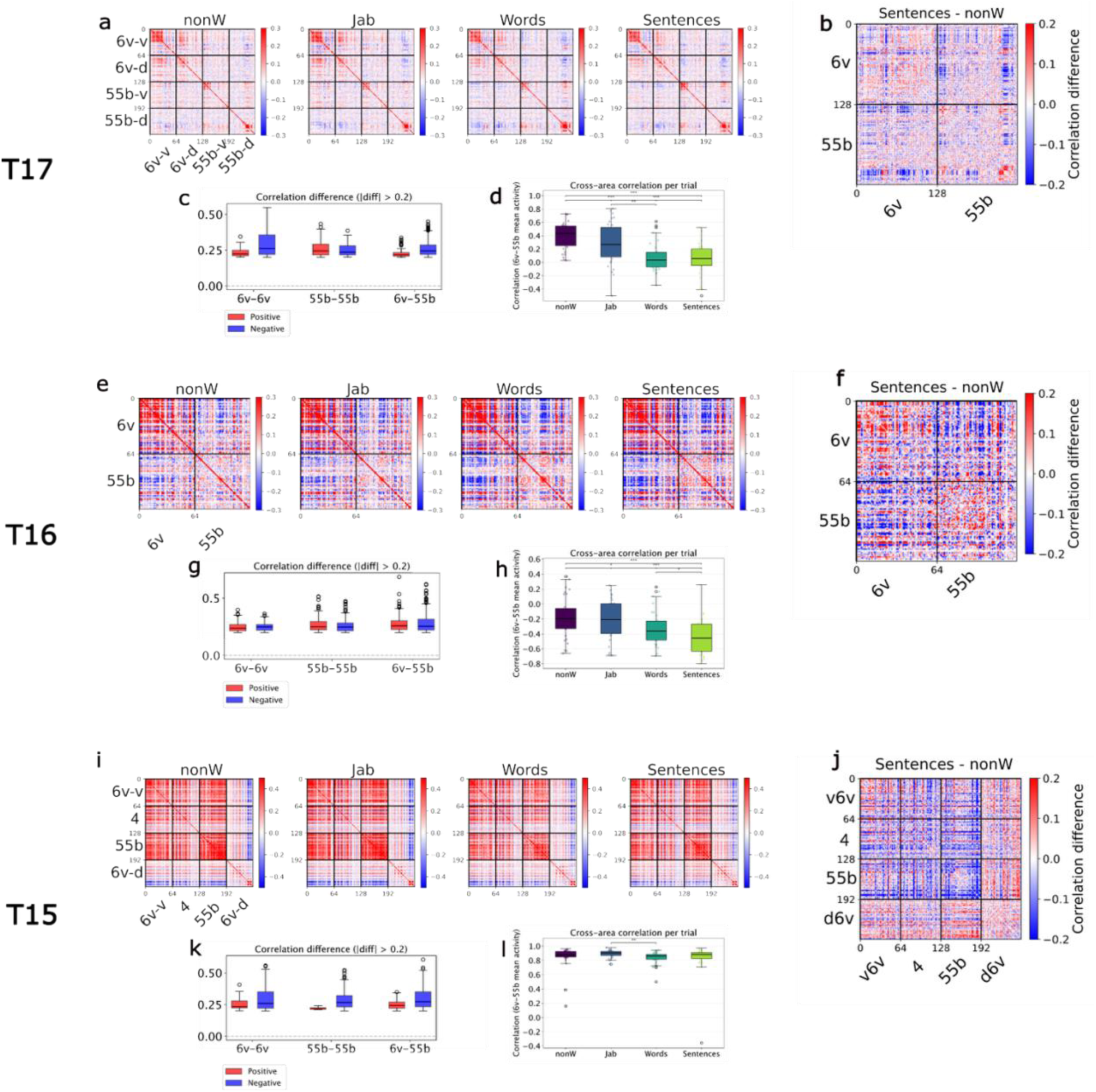
Correlation analysis within and across areas 6v and 55b. **a,e,i)** Pairwise correlation matrices per participant and condition. The correlation matrices were calculated per trial and averaged across trials for each condition. **b,f,j)** The difference between the sentences and nonwords pairwise correlation matrices. **c,g,k)** Box plots showing the distribution of correlation difference values (sentences minus nonwords, as shown in Figure 2g– i) for channel pairs where the absolute difference exceeds 0.2. Values are separated by sign of the difference (positive: sentences > nonwords; negative: nonwords > sentences) and by channel pair type (within 6v, within 55b, or between 6v and 55b). **d,h,i)** Box plots showing the distribution across trials of the mean correlation between 55b and 6v channel pairs, per condition.

**Extended Data Figure 4:**
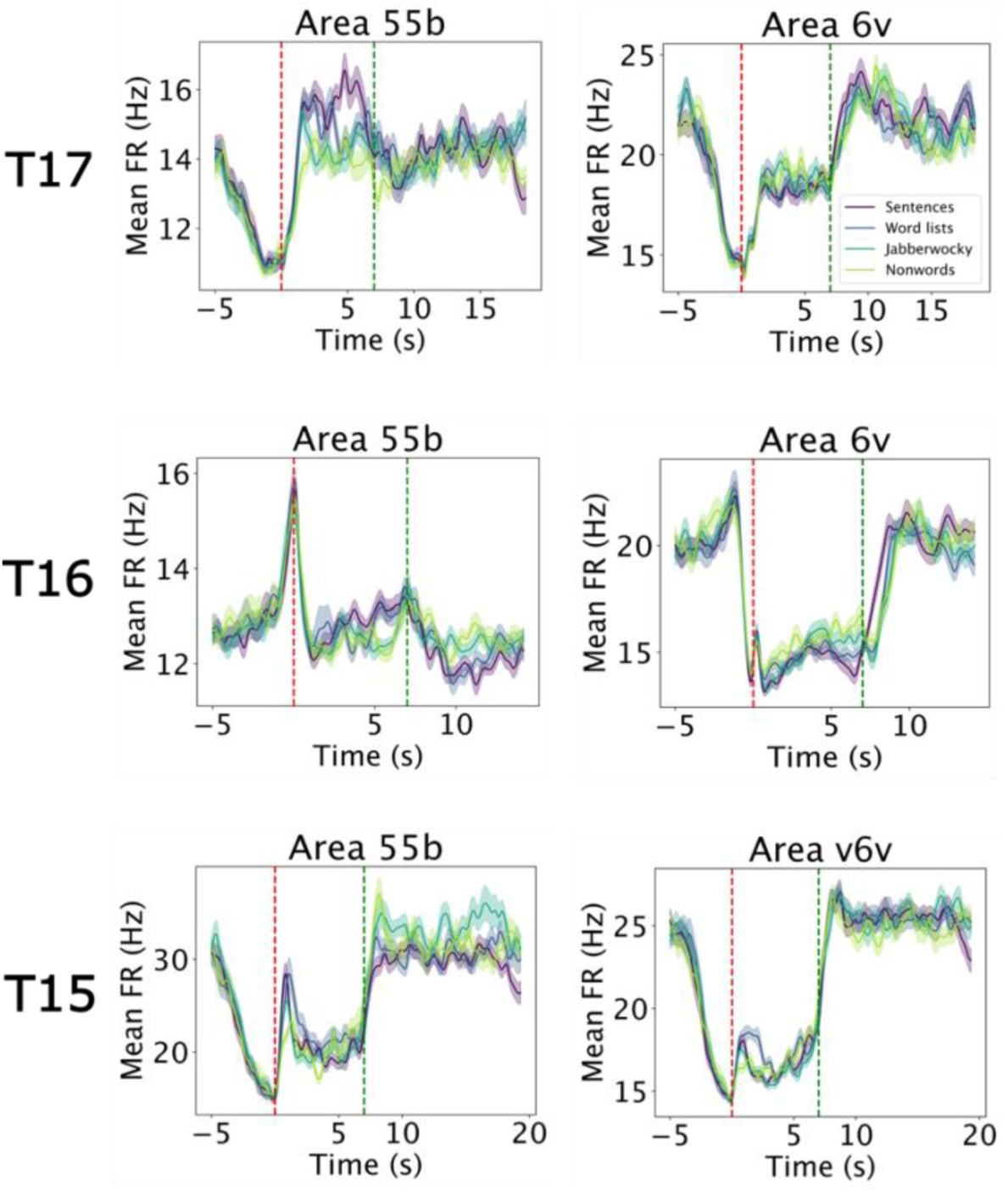
Similar to Figure 1e in the main text, but starting from trial onset (t=0). Averaged PSTH for area 55b (left) and area 6v (right) of each participant. Firing rates were averaged over all 128, 64 and 64 channels in each area for participants T17, T16 and T15 respectively. This figure shows the waveform for the whole trial where 0 is trial onset and dashed vertical green line denotes the average Go cue time at 7s (in the task Go cue time was at 7±0.5 s following trial onset).

**Extended Data Figure 5:**
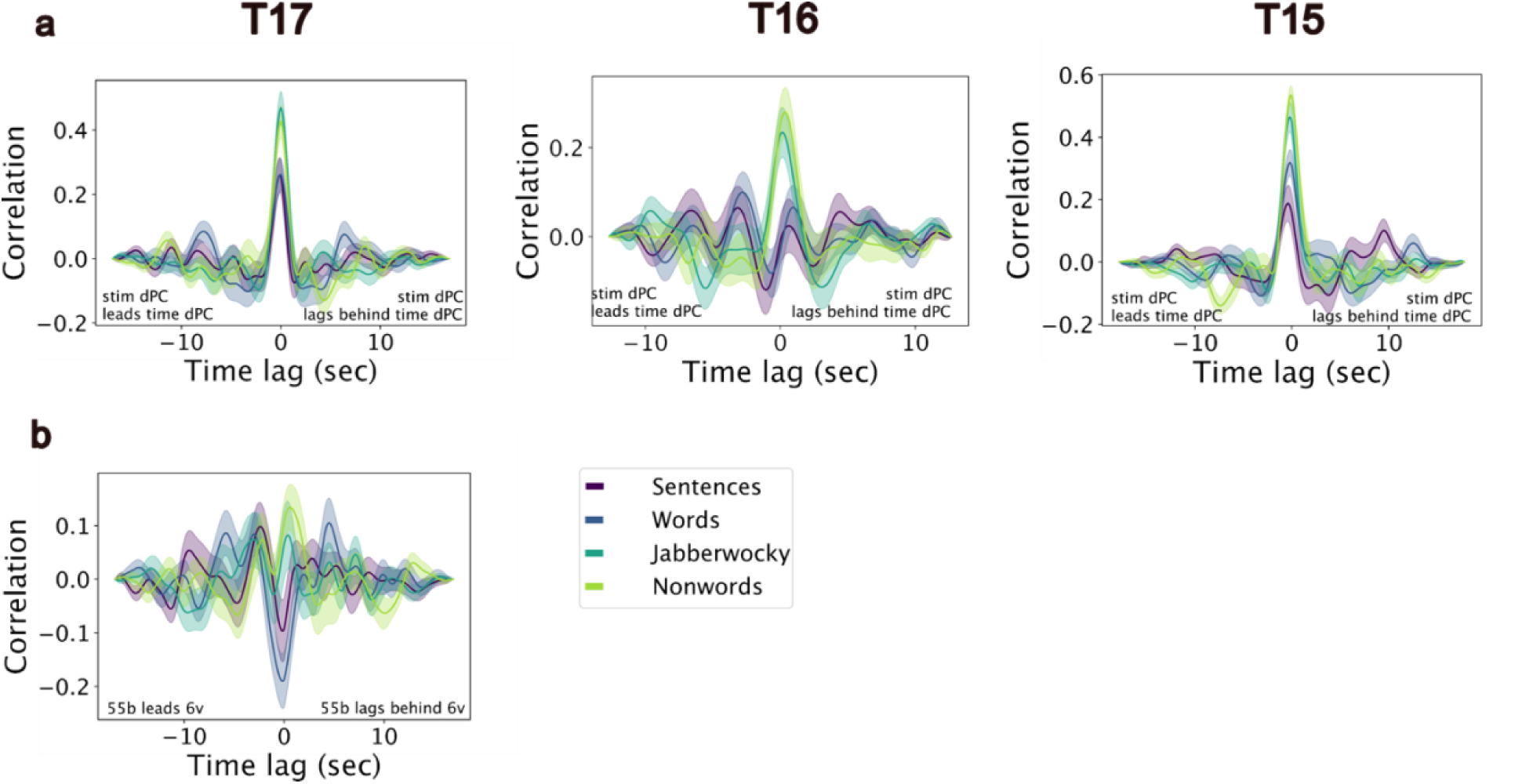
Time-lagged cross correlation of condition-centered data. Similar to Fig. 3d,e in the main text, but correlations are calculated after subtracting the mean across trials per condition. **a)** Cross-correlation in time between the first stimulus dPCA component and the first time dPCA component, after the mean across trials was subtracted for each condition in both components. The same dPCA components were used here as for Fig. 3. **b)** Cross-correlation in time between T17’s 55b channel-averaged signal and 6v channel-averaged signal, after subtracting the mean across trials for each condition.

**Extended Data Figure 6:**
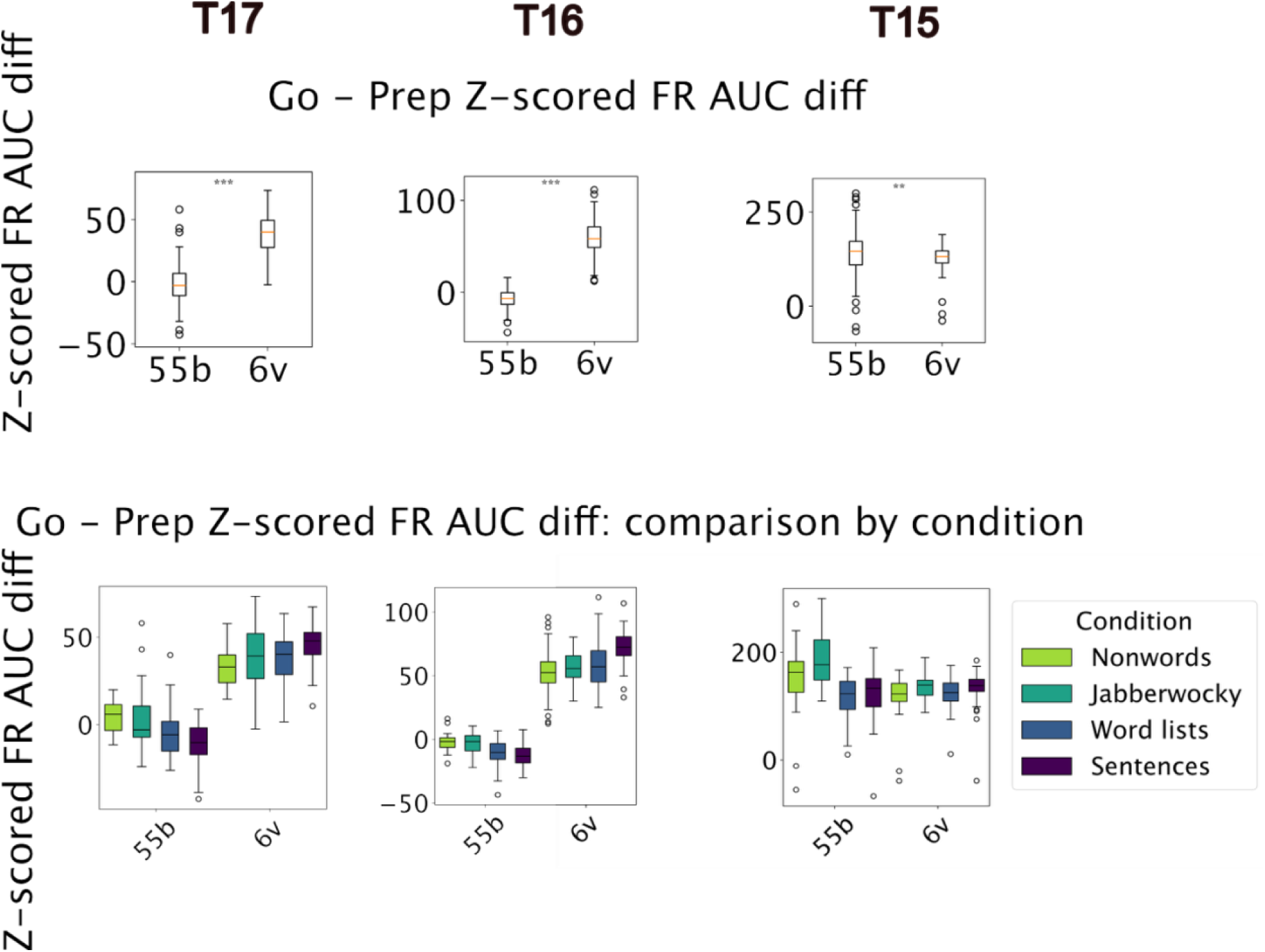
Comparison of firing rates (FR) between Go and preparatory periods. The z-scored area under the firing rate curve (FR AUC; as defined in Fig. 1 in the main text) during the Preparatory period was subtracted from the AUC during the Go period on a trial-by-trial basis. The resulting AUC difference (Go − Prep) is shown for each brain area. **(Top)** The AUC difference is shown with all trials pooled across stimulus conditions. Asterisks denote significant differences between brain areas (*: p<0.05, **: p<0.01, *** p<0.001, Student’s t- test). **(Bottom)** The AUC difference is shown separately for each stimulus condition.

**Extended Data Figure 7:**
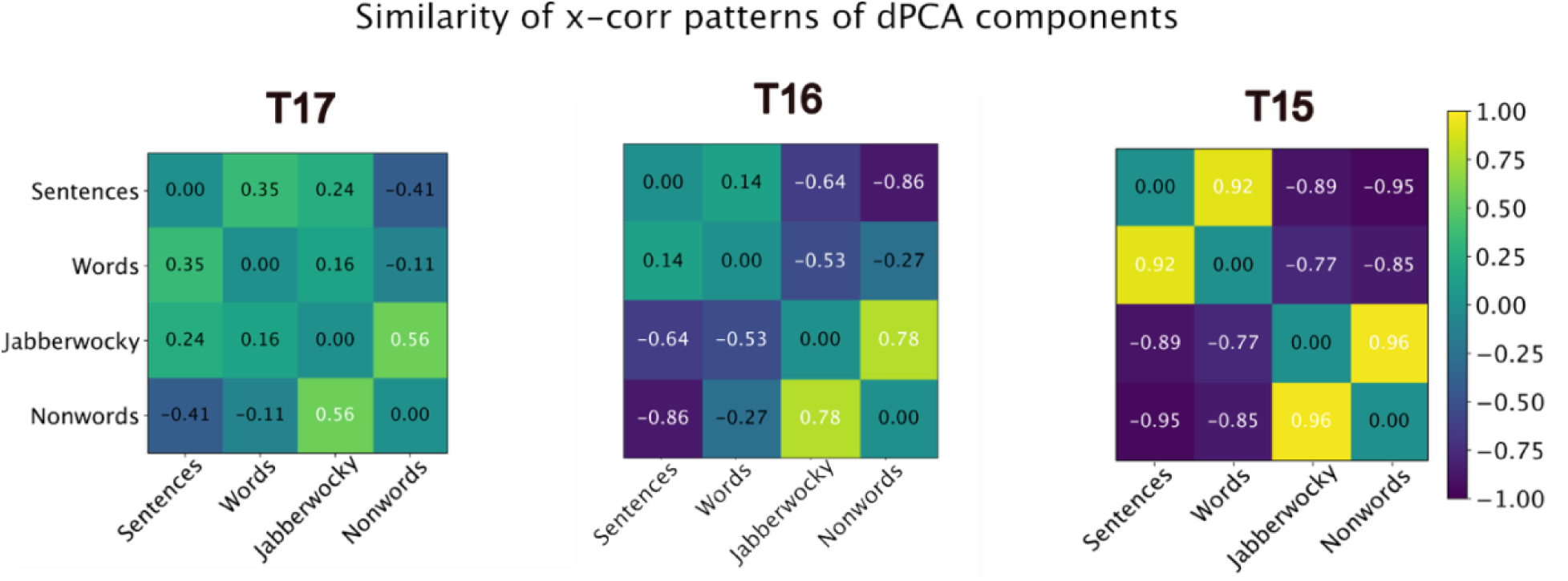
Similarity between time-lagged cross-correlation patterns across conditions. The similarity between the time-lagged cross-correlation patterns of the stimulus and time dPCs was calculated between each pair of the four stimulus conditions shown in Fig. 3d in the main text, using Pearson’s correlation coefficient (r). Correlation values are shown both as text within the matrices and by the heatmap color bar.

## Extended information 2: Example sentences from Switchboard dataset clusters

### T17’s clusters

Cluster 0 example sentences:

- I think that’s not very smart.

- We are not real consistent with this.

- I think not normal setting.

- So I figured it wasn’t true.

- It’s kind of a depressing topic.

- You can’t enjoy it.

- It’s really hard to.

- Leaned the wrong way.

- That’s very hard to do.

- Really a rough time you know.

Cluster 1 example sentences:

- What do we recycle?

- If you get pleats.

- Except you get to taste the product instead.

- What is the other one called?

- They’re both declawed.

- Tried to break that off.

- Get the recipe from them.

- I guess what I’m trying to get at is.

- There is a receptacle.

- Rehabilitate you.

Cluster 2 example sentences:

- I was in education and in administration.

- I went back to school.

- Well we have one through work.

- I work in the waiver.

- For a checkup or whatever.

- They have one of the clerks up there.

- The question was talking about the juries.

- So you did buy a pattern.

- About the same I dress for school.

- You could mail it in.

Cluster 3 example sentences:

- It’s really nice.

- They’ve got a nice one.

- It’s a real good concept.

- It’s really cute.

- That sounds real good.

- It’s something we like.

- I really enjoy that.

- I’m really enjoying it so far.

- I think it’s great.

- It’s worth doing it.

Cluster 4 example sentences:

- That’s just what it is.

- They say that it is.

- That’s what I mean.

- That’s pretty much what it is.

- That’s what they say.

- That’s kind of the same thing for me.

- That’s the way I feel.

- We do, as a matter of fact.

- Well I’m definitely for it.

- Yeah I mean that.

Cluster 5 example sentences:

- It seems like we pay the state.

- It’s an association fee.

- It’s aking profits.

- Owning on them and making payments on them.

- We take a certain portion of your paycheck.

- The Treasury view would be that.

- It was paid for.

- The time frame which they get paid.

- You do have government money to fund.

- And money is their main goal.

Cluster 6 example sentences:

- Go to a trip through the Indonesian.

- I called the minute the sun went down.

- Knox is just over the hill here.

- Country western day or something like that one.

- We got a chance to make some noise this year.

- I saw something the other night about it.

- I remember one time up in Nashville.

- After that season, Alex had a decision to make.

- We’re going to buy a Devon Rex.

- I’ve got a Los Angeles map.

### T15’s clusters

Cluster 0 example sentences:

- That does not seem quite fair.

- That does not seem quite fair.

- We are not real consistent with this.

- Certainly the purpose is not very ethical.

- Saying this does not make it right.

- No, I do not feel comfortable.

- It doesn’t fit the circumstance anymore.

- It isn’t a very wise decision.

- No, I do not feel very comfortable.

- It seems that having gone through that is not helpful.

Cluster 1 example sentences:

- I’m making some food for us.

- I will cook at least something to eat.

- We often like to eat local food.

- I’d love to join you to eat.

- I like to bake off and on.

- I will order some food for everyone here.

- I usually love to eat there.

- How do you usually eat it?

- That’s why I like some diversity in my breakfast food.

- This couple is known to cook good food.

Cluster 2 example sentences:

- Someone will give you a call next week.

- I can give you a call next week.

- As you know, we will go there next week.

- We should go there next week.

- Maybe I will use it next week.

- And then we rent them for a week.

- It came out in the first week of the year.

- It came out in the first week of the year.

- I can see you next week.

- Business was established last week.

Cluster 3 example sentences:

- The money speaks for us.

- It seems like we pay the state.

- How much do I have to pay you?

- When they recognize that it’s their tax money.

- Then they’ll pay you a certain amount of money.

- A certain amount of their salary would go to it.

- Bring that paid tax report.

- We can deduct that expense from our income tax.

- A lot of it goes down to just money.

- The time frame which they get paid.

Cluster 4 example sentences:

- My house has a living room.

- We just moved into this house.

- I use them both here at work and at home.

- She lives in the house right next to me.

- You should see our house.

- How are things at home?

- We moved into the apartment and got cable.

- Trying to find someone at home.

- I work out of my home.

- Just in house programs to do what they needed.

Cluster 5 example sentences:

- Blizzards now have a new visual effect.

- It was weird because we had snow.

- We lived up north long enough to appreciate the cold.

- Winter weather is kind of a problem around here.

- North side is getting very cold lately.

- This is unusual for the people in the north.

- This past winter was very cold.

- How many states in America get a lot of snow?

- ice hairs blooming coming opinion take

- You’ve had more freezing this Winter.

Cluster 6 example sentences:

- Can you play with my dog for a bit?

- We have a dog too so that adds to it.

- My dog will play with his food.

- Even if I have my dog with me.

- I have seen him taking care of the dog.

- I want to take my dog for a walk.

- I like miniature poodles and other small dogs.

- You like pets and things.

- We’ve got so many when he was a puppy.

- Well at the present time I have two dogs.

Cluster 7 example sentences:

- A couple of children and we have to.

- Someplace for the children to be.

- Let’s make a program for kids.

- Once they have children over here.

- To where the children are.

- The event was really oriented toward little kids.

- If young people had to do it.

- Noticed that when I was a kid.

- My children brought it to my attention.

- My children brought it to my attention.

Cluster 8 example sentences:

- They sent us a little booklet to read.

- I can walk to get the books.

- I guess I need to put it on paper and turn it in.

- I read those little slips that come in your mail.

- Pay for paper to recycle.

- Make sure you read it again.

- She wrote my name on a list.

- I sold a bunch of books that I read.

- He’s got a couple of books out.

- Could you bring that paper with you?

Cluster 9 example sentences:

- It seems to be really good for them.

- I think it is a wonderful place.

- It makes it very convenient for us.

- Those are the ones I really enjoy.

- I have fun doing it.

- Of course it’s real convenient for you.

- But it was great pleasure while we did it.

- So it’s been real fun here to see.

- It’s really nice to see them.

- I think it’s pretty exciting that they can do that.

